# Investigating a role for *PUB17* and *PUB16* in the self-incompatibility pathway in transgenic *Arabidopsis thaliana*

**DOI:** 10.1101/2023.10.24.563783

**Authors:** Paula K.S. Beronilla, Daphne R. Goring

**Affiliations:** Department of Cell & Systems Biology, University of Toronto, Toronto, Canada M5S 3B2; Centre for the Analysis of Genome Evolution & Function, University of Toronto, Toronto, Canada M5S 3B2

**Author notes:** Corresponding Author: Daphne Goring.

**Keywords:** *Arabidopsis*, *Brassica*, self-incompatibility, SRK, ARC1, PUB17, PUB16

## Abstract

In Brassicaceae self-incompatibility (SI), self-pollen rejection is initiated by the *S-*haplotype specific interactions between the pollen SCR/SP11 ligand and the stigma S Receptor kinase (SRK). In *Brassica* SI, a member of the Plant U-Box (PUB) E3 ubiquitin ligases, ARC1, is then activated by SRK in this stigma and cellular events downstream of this cause SI pollen rejection by inhibiting pollen hydration and pollen tube growth. During the transition to selfing, *Arabidopsis thaliana* lost the SI components, *SCR, SRK*, and *ARC1*. However, this trait can be reintroduced into *A. thaliana* by adding back functional copies of these genes from closely related SI species. Both *SCR* and *SRK* are required for this, though the degree of SI pollen rejection varies between accessions, and *ARC1* is not always needed to produce a strong SI response. For *A. thaliana* C24, only transforming with *A. lyrata SCR* and *SRK* confers a strong SI trait, and so here we investigated if ARC1-related PUBs were involved in the SI pathway. Two close ARC1 paralogs, *PUB17* and *PUB16*, were selected, and CRISPR/Cas9 technology was used to generate *pub17* and *pub16* mutations in the C24 accession. These mutants were then crossed into a transgenic *A. thaliana* SI-C24 line and their potential impact on SI pollen rejection was investigated. Overall, we did not observe any significant differences to implicate *PUB17* and *PUB16* functioning in the transgenic *A. thaliana* SI-C24 stigma to reject SI pollen.

## 1 INTRODUCTION

In the Brassicaceae, the dry stigma employs a strong selectivity for pollen as it determines the fate of the pollen in contact through the activation of intracellular events to facilitate pollen hydration (Dickinson, 1995). This is followed by the germination of the pollen tube which penetrates the stigmatic cell wall and travels down the reproductive tract until it reaches an unfertilized ovule for fertilization (Hafidh and Honys, 2021, Robichaux and Wallace, 2021). In self-incompatible (SI) Brassicaeae species, the stigma blocks self-incompatible pollen by preventing pollen hydration and germination (Abhinandan *et al*., 2022, Goring *et al*., 2023). Brassicaceae SI is genetically controlled by two *S*-locus linked genes encoding the pollen S Cysteine-Rich/ S-locus Protein 11 (SCR/SP11) peptide (Schopfer *et al*., 1999, Takayama *et al*., 2000, Shiba *et al*., 2001) and the stigma *S* Receptor Kinase (SRK) (Stein *et al*., 1991, Goring and Rothstein, 1992, Takasaki *et al*., 2000, Silva *et al*., 2001). Both genes are highly polymorphic and have specific allelic combinations (*S*-haplotypes) that determine SI pollen recognition. Following *S*-haplotype specific receptor-ligand interactions between the stigma SRK and the pollen SCR, SRK activates the SI signalling pathway in the stigmatic papilla to reject the SI pollen. SCR and SRK have been primarily studied in *Brassica* and *Arabidopsis* species, but the signaling events and mechanisms underlying the SI pathway have been best characterized in *Brassica* species (Abhinandan *et al*., 2022, Goring *et al*., 2023).

There are three *Brassica* SRK interacting proteins that have been shown to function in the *Brassica* SI pathway: ARM-Repeat-Containing 1 (ARC1; Gu *et al*., 1998, Stone *et al*., 1999), M Locus Protein Kinase (MLPK; Murase *et al*., 2004, Kakita *et al*., 2007), and the FERONIA (FER) receptor kinase (Zhang *et al*., 2021, Huang *et al*., 2023). *Brassica napus* ARC1 is a Plant U-box (PUB) E3 ubiquitin ligase (Gu *et al*., 1998, Azevedo *et al*., 2001, Stone *et al*., 2003) and was shown to be a key player in the stigma SI pathway through the partial and complete breakdown of SI pollen rejection with RNAi knockdowns and CRISPR/Cas9 mutations, respectively (Stone *et al*., 1999, Abhinandan *et al*., 2023, Huang *et al*., 2023). In the SI pathway, *B. napus* ARC1 confers pollen rejection by targeting compatibility factors in the stigma for ubiquitination and proteasomal degradation. The identified targets include Glyoxylase 1 (GLO1; Sankaranarayanan *et al*., 2015), the EXO70A1 exocyst subunit (Samuel *et al*., 2009, Safavian and Goring, 2013), and phospholipase D α1 (PLD1; Scandola and Samuel, 2019). The degradation of these factors through ARC1 leads to SI pollen rejection by disrupting cellular responses in the stigmatic papillae that are needed for compatible pollen (cellular homeostasis, exocytosis, and membrane remodelling; Abhinandan *et al*., 2022). Recently, *B. rapa* FER was implicated as another signaling branch downstream of SRK (Zhang *et al*., 2021, Huang *et al*., 2023). It is proposed that SRK interacts with FER which then activates NADPH oxidases causing the accumulation of reactive oxygen species (ROS) to inhibitory levels (Zhang *et al*., 2021, Huang *et al*., 2023). The SRK-activated ARC1 and FER pathways function in parallel and disrupting either causes a breakdown in SI pollen rejection.

In *Arabidopsis* species, identifying SRK interacting proteins has proven to be more elusive, but some downstream cellular responses following SI pollen perception have been uncovered (Safavian and Goring, 2013, Indriolo *et al*., 2014, Iwano *et al*., 2015, Macgregor and Goring, 2022, Macgregor *et al*., 2022). While *A. thaliana* carries pseudogenized *SCR, SRK*, and *ARC1* genes, the SI trait can be reinstated using transgenes from SI *Arabidopsis* species such as *A. lyrata* and *A. halleri* (Nasrallah *et al*., 2004, Boggs *et al*., 2009b, Indriolo *et al*., 2014, Iwano *et al*., 2015, Zhang *et al*., 2019, Rozier *et al*., 2020). However, accession differences were uncovered in these studies where, for example, the Al*SCR* and Al*SRK* transgenes successfully conferred a strong SI phenotype in C24 but not in Col-0 (Nasrallah *et al*., 2004, Boggs *et al*., 2009b, Indriolo *et al*., 2014, Iwano *et al*., 2015). This variation was found to be caused by RNA-induced silencing of the Al*SRK* transgene by small RNAs originating from inverted repeats in the expressed *A. thaliana* Col-0 *SRKA* pseudogene (Fujii *et al*., 2020). We previously found that *ARC1* is required for the SI trait in *A. lyrata* and adding Al*ARC1* along with the Al*SCR* and Al*SRK* transgenes into *A. thaliana* Col-0 also restored the SI phenotype (Indriolo *et al*., 2012, Indriolo *et al*., 2014). The interaction of AlARC1 with SRK was found to be conserved, and interestingly a close ARC1 paralog, PUB17, interacted with SRK as well (Indriolo and Goring, 2016). This raised the question of whether ARC1 paralogs could also be functioning in *Arabidopsis* SI. Here, using a transgenic *A. thaliana* Al*SCR* Al*SRK* C24 line that displays a strong SI phenotype (Iwano *et al*., 2015), we investigated the involvement of two close ARC1 paralogs, PUB16 and PUB17, in the transgenic *A. thaliana* SI pathway.

## 2 RESULTS

### Generation of *pub16* and *pub17* mutations in the C24 accession

ARC1 belongs to a subgroup of PUB E3 ligases that have the domain organization of a N-terminal UND domain, followed by a U-box domain and C-terminal ARM domain (Mudgil *et al*., 2004, Samuel *et al*., 2008, Trenner *et al*., 2022). A phylogeny of AlARC1 with UND-PUBs from *A. lyrata* and *A. thaliana* showed that the closest paralogs to AlARC1 are PUB17 and PUB16 (Figure 1; Indriolo *et al*., 2012). Since AlPUB17 was also previously found to interact with AlSRK_1_ (Indriolo and Goring, 2016), PUB17 and PUB16 were selected to investigate if they functioned in the transgenic *A. thaliana* SI pathway. The transgenic *A. thaliana* Al*SCR* Al*SRK* C24 line (SI-C24 #15-1) was selected for this study as it displays a strong and stable SI phenotype (Iwano *et al*., 2015, Macgregor *et al*., 2022). *PUB17* and *PUB16* are expressed across a broad range of tissues in *A. thaliana* including the stigma (Klepikova *et al*., 2016), and the expression of *PUB17* and *PUB16* in pistil tissue was confirmed by RT-PCR (Figure 2a).

**FIGURE 1.**
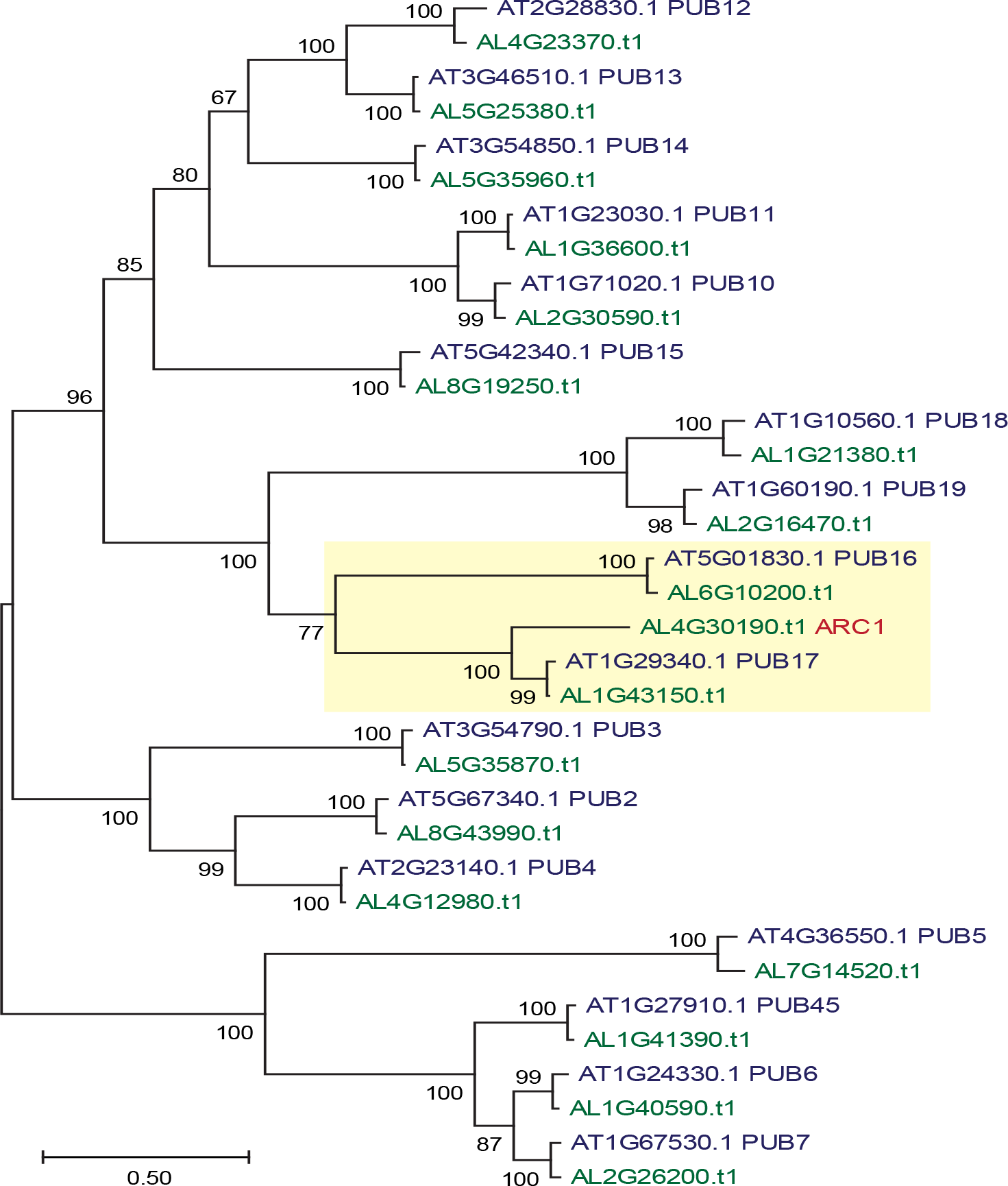
Phylogenetic analysis of predicted UND-PUB proteins in *Arabidopsis thaliana* and *Arabidopsis lyrata*. The amino acid sequences for *A. thaliana* UND-PUBs are shown in blue and *A. lyrata* UND-PUBs are in green (the *A. lyrata* ARC1 name is in red).

**FIGURE 2.**
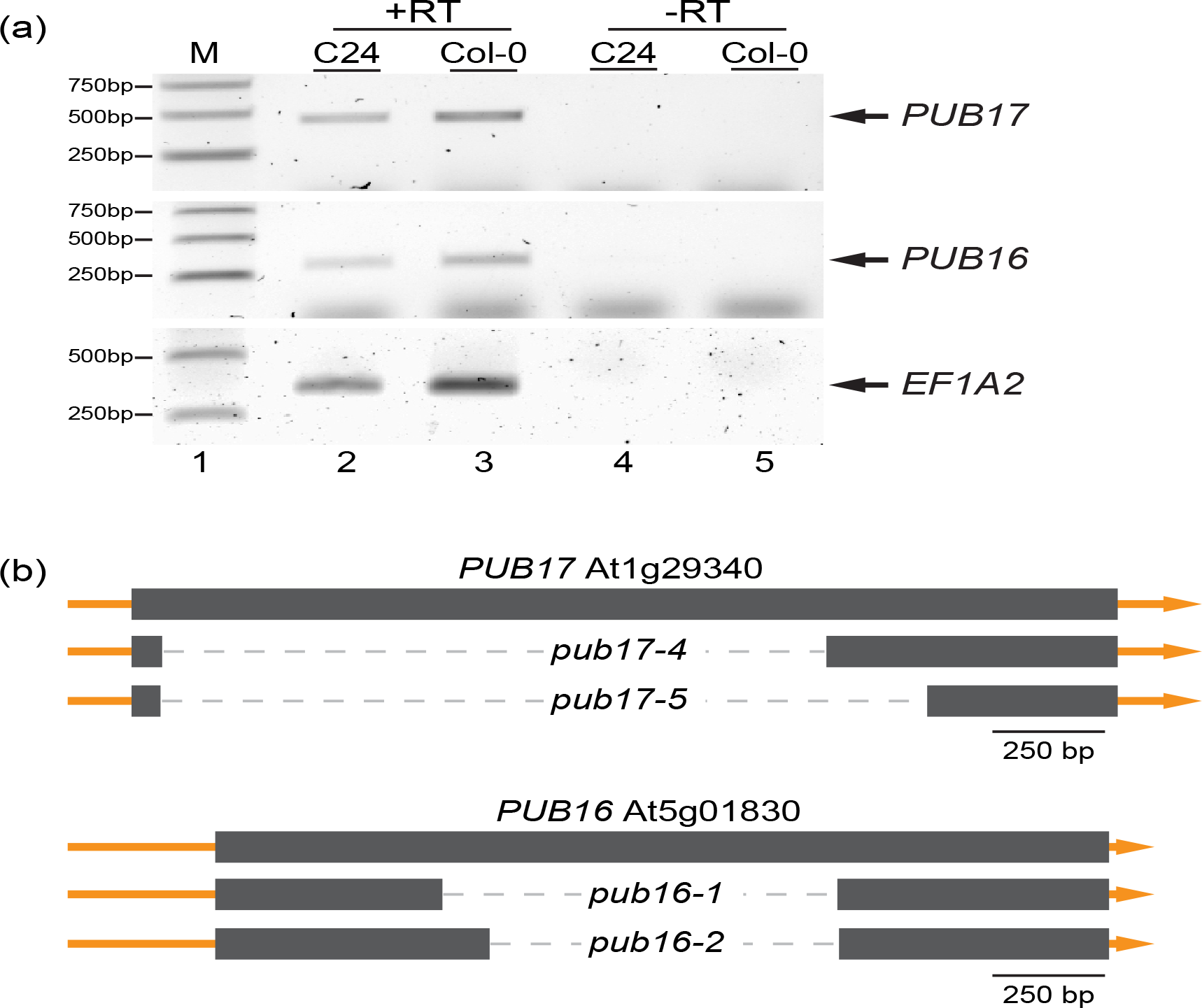
Gene expression of PUB17 and PUB16 in *A. thaliana*.*(a)*RT-PCR analysis of *PUB17* and *PUB16* expression in pistil tissues from *A. thaliana* Col-0 and C24. cDNA was synthesized from RNA extracted from the top half of pistils (including the stigmas), and RT-PCR was conducted to confirm expression of *PUB17* and *PUB16* in both accessions. *EF1A2* was used as a positive control. *PUB17*= 467 bp, *PUB16*= 305 bp, *EF1A2*= 355bp.*(b)*Schematic representation of CRISPR/Cas9 deletion mutants for *PUB17* and *PUB16*. Both genes are encoded by a single exon, and dashed lines represent deletions in the mutant alleles. Scale bars = 250 bp.

Since *pub17* and *pub16* mutants were not available in the C24 accession, we generated individual mutants using CRISPR/Cas9 vectors targeting each gene with two sgRNAs designed to generate deletion mutations (Figure 2b). Wildtype *A. thaliana* C24 was transformed with these vectors, and transformed T1 plants carrying deletion mutations were identified by PCR. Homozygous mutants were detected in the T2 generation, and all were wildtype in appearance (*pub17-4, pub17-5, pub16-1, pub16-2*). These *pub* mutants were then crossed individually into the SI-C24 line, and SI-C24 plants that were homozygous for either the *pub17* or *pub16* mutations were identified in the T2 generation for further analysis. Finally, to investigate any potential redundancy between *PUB17* and *PUB16*, the *pub16-1* mutant was crossed with the SI-C24 *pub17-5* mutant line to generate the SI-C24 *pub16-1 pub17-5* double homozygous mutant line.

### SI pollen hydration on SI-C24 *pub* mutant stigmas

The rejection of SI pollen by the stigma is quite rapid and can been seen in reduced SI pollen hydration. Thus, we tested the impact of the *pub* mutations on the transgenic *A. thaliana* SI pathway by conducting pollen hydration assays (Figure 3). For this assay, SI-C24 pollen was used to pollinate wildtype and mutant stigmas, and pollen diameters were measured as a proxy for pollen hydration at 0 and 10-min post-pollination (Lee *et al*., 2020). When SI-C24 pollen was used to pollinate compatible C24 stigmas, there was a large increase in pollen diameter due to compatible pollen acceptance (Figure 3a-c). Large increases in SI-C24 pollen diameter were also observed on the *pub17* and *pub16* mutant stigmas, and the average pollen diameter at 10-min post-pollination was not significantly different from that seen on the C24 stigmas (Figure 3a-b). These results indicate that the *pub17* and *pub16* mutations in the C24 background did not disrupt any early compatible post-pollination responses (Figure 3a-b). Next, we tested the potential impact of the *pub17* and *pub16* mutations on the SI response in the transgenic *A. thaliana* stigma. SI-C24 pollen placed on SI-C24 stigmas resulted in only a small increase in pollen diameter due to SI pollen rejection (Figure 3a-c). Similarly, when SI-C24 pollen was used to pollinate stigmas from the SI-C24 *pub17* and SI-C24 *pub16* lines, there were small increases in pollen diameter and the average pollen diameters at 10-min post-pollination were not significantly different to that for SI-C24 stigmas (Figure 3a-b). These results indicate that the individual *pub17* and *pub16* mutations in the SI-C24 background did not have any noticeable impact on the early SI pollen rejection responses. Finally, to address whether *PUB17* and *PUB16* may have redundant functions in the rejection of SI pollen, pollen hydration assays were conducted on SI-C24 *pub16-1 pub17-5* mutant stigmas (Figure 3c). SI-C24 pollen placed on the SI-C24 *pub16-1 pub17-5* mutant stigmas exhibited a small increase in SI pollen hydration similar to that observed for SI-C24 stigmas, which indicates that SI pollen hydration was unaffected by the mutation of both *PUB17* and *PUB16* in the SI-C24 stigma. As a control, C24 pollen was used to pollinate the SI-C24 *pub16-1 pub17-5* mutant stigmas, and these pollen grains hydrated normally confirming that the combined *pub16-1 pub17-5* mutations did not impact compatible pollen hydration as well (Figure 3d). The collective results from the pollen hydration assay suggest that *PUB17* and *PUB16* are not required in the transgenic *A. thaliana* SI-C24 stigma for the rejection of SI-C24 pollen at the pollen hydration stage.

**FIGURE 3.**
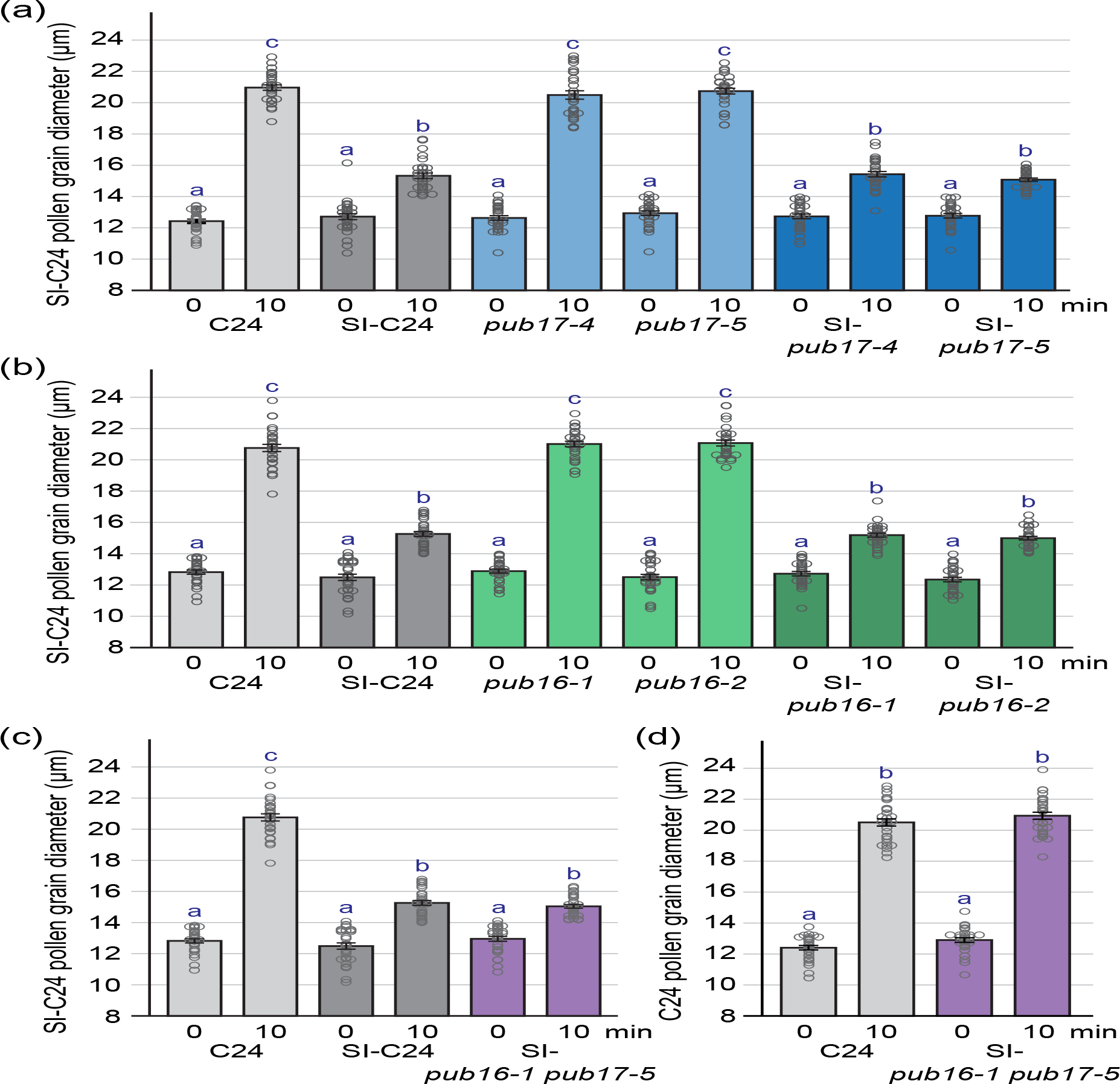
SI-C24 pollen hydration assays on SI-C24 and SI-C24 *pub* mutant stigmas. (a-c) SI-C24 pollen hydration on stigmas from (a) C24, SI-C24, *pub17* and SI-C24 *pub17* mutants; (b) C24, SI-C24, *pub16* and SI-C24 *pub16* mutants; and (c) C24, SI-C24 and the SI-C24 *pub16-1 pub17-5* mutant. (d) C24 pollen hydration on stigmas from C24 and the SI-C24 *pub16-1 pub17-5* mutant. Pollen diameters were measured at 0- and 10-min post-pollination. Data are shown as bar graphs of the means ± SE with all the data points displayed. n = 30 pollen grains per line. Letters represent statistically significant groupings of P < 0.05 based on a one-way ANOVA with a Tukey-HSD post-hoc test.

### SI pollen tube growth and seed set in the SI-C24 *pub* mutants

The majority of SI pollen grains are rejected by blocking pollen hydration, but some do successfully hydrate and produce a pollen tube that grows into the pistil. As a result, we wanted to further investigate if the loss of *PUB17* and *PUB16* had any impact on the number of pollen tubes that successfully overcame the SI stigma barrier. Control and mutant pistils were pollinated with SI-C24 pollen and collected at 24-h post-pollination for aniline blue staining to visualize the pollen tubes (Figure 4). A control compatible pollination of C24 pistils with SI-C24 pollen showed abundant pollen tubes growing through the C24 pistils signifying pollen acceptance (Figure 4a, b). In contrast, the control SI pollination of SI-C24 pistils with SI-C24 pollen resulted in an absence of pollen tubes growing through these pistils indicating pollen rejection (Figure 4c-d). Compatible pollinations of the *pub17* and *pub16* mutants with SI-C24 pollen resulted in abundant pollen tube growth through the pistils (Figure 4e-f, i-j). As well, compatible pollinations of the SI-C24 *pub16-1 pub17-5* mutant pistils with C24 pollen resulted in many pollen tubes growing through the pistils (Figure 4m-n). There were no observable differences in the abundance of pollen tubes growing through the *pub* mutant pistils in comparison to C24 signifying that pollen tube growth was not affected in these mutants.

**FIGURE 4.**
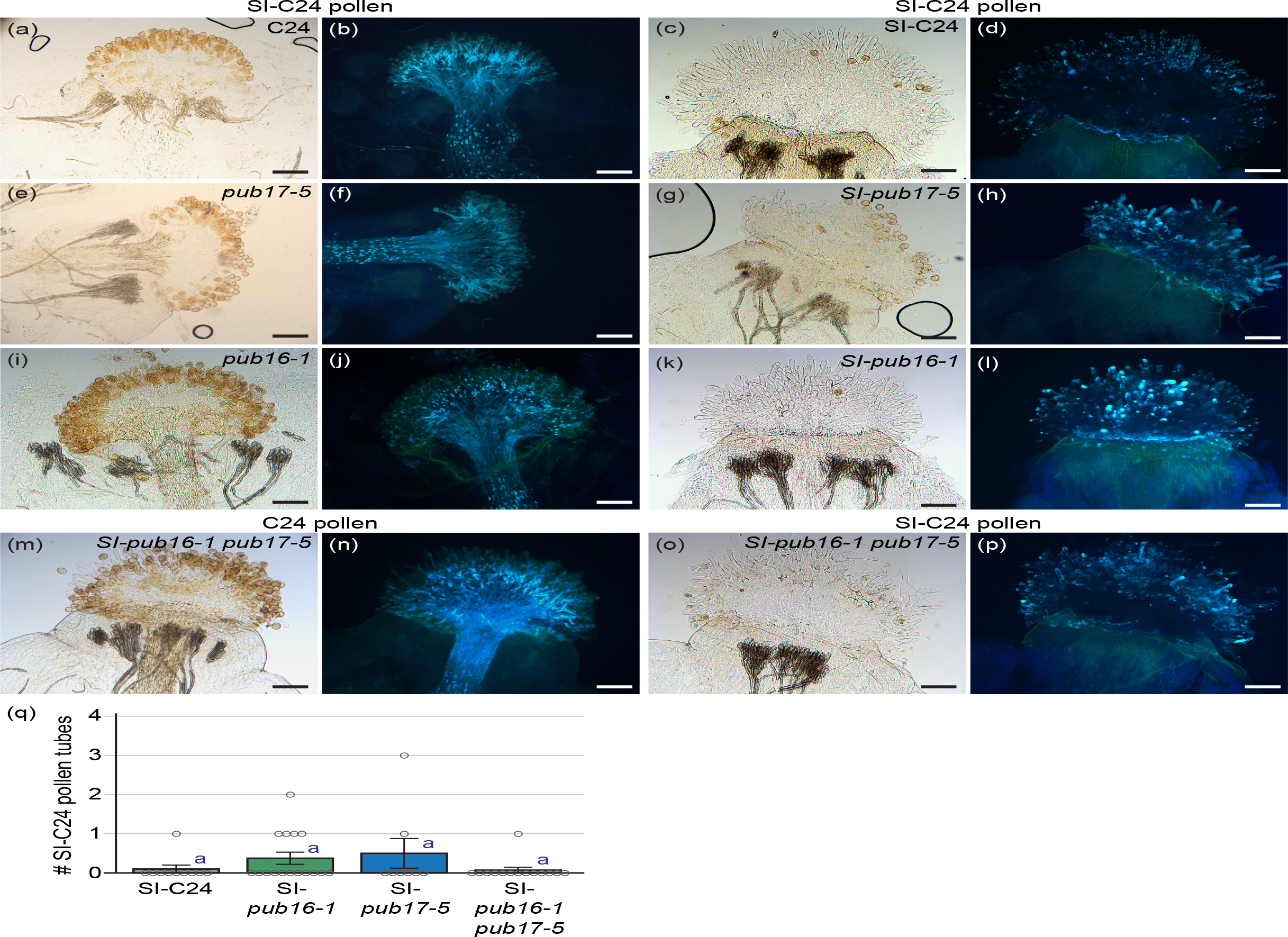
Aniline blue images of SI-C24 pollinated SI-C24 and SI-C24 *pub* mutant pistils.(a-p) Representative images of pistils collected at 24-h post-pollination for aniline blue staining. The pistils were pollinated with SI-C24 pollen, except for (m-n) which were pollinated with C24 pollen. The genotypes of the pistils are indicated above, and brightfield (left) and aniline blue images (right) are shown for each sample. Pollen tubes growing through the pistil are indicative of pollen acceptance. Scale bars = 100 µm.(q) Bar graph showing the mean number of pollen tubes/pistil at 24-h post-pollination. Data are shown as mean ± SE with all the data points displayed. n = 8-16 pistils per line. Letters represent statistically significant groupings of P < 0.05 based on a one-way ANOVA with a Tukey-HSD post-hoc test.

When SI-C24 *pub17-5* and SI-C24 *pub16-1* mutants were pollinated with SI-C24 pollen, pollen tubes could not be observed in the pistils (Figure 4g-h, k-l). Similarly, this phenotype was observed in the SI-C24 *pub16-1 pub17-5* mutant pistils pollinated with SI-C24 pollen (Figure 4o-p). To verify that there was no impact on SI-C24 pollen rejection at the level of pollen tube growth, the number of pollen tubes in the style were counted in the SI-C24 and SI-C24 *pub* mutant pistils (Figure 4q). While there was a slight trend of more pollen tubes growing through the SI-C24 *pub17-5* and SI-C24 *pub16-1* mutant pistils, it was not significantly different to that observed for the SI-C24 pistils. Thus, these data indicate that the loss of *PUB17* and *PUB16* function in the transgenic *A. thaliana* SI-C24 pistil did not impact the rejection of SI-C24 pollen tube growth.

While the *pub* mutations did not disrupt SI-C24 pollen rejection at the hydration and pollen tube growth stages, we wanted to see if there would be any impact on the final stage of seed production. The SI-C24 *pub* mutants were pollinated with C24 and SI-C24 pollen, the number of seeds/siliques were counted (Figure 6). With compatible C24 pollen, abundant seeds were produced for all samples and SI-C24 *pub* mutants did not show any significant differences to the SI-C24 control (Figure 6). Interesntigly with SI-C24 pollen, there was the same trend as seen with the number of pollen tubes (Figure 4q) where the SI-C24 *pub17-5* and SI-C24 *pub16-1* mutant siliques has a slight increase in the number of seeds per silique, but they were not significantly different to the SI-C24 siliques (Figure 5). Overall, these results indicate that the loss of *PUB17* and *PUB16* in the transgenic *A. thaliana* SI-C24 pistil did not impact seed yield with either compatible C24 or SI-C24 pollinations.

**FIGURE 5.**
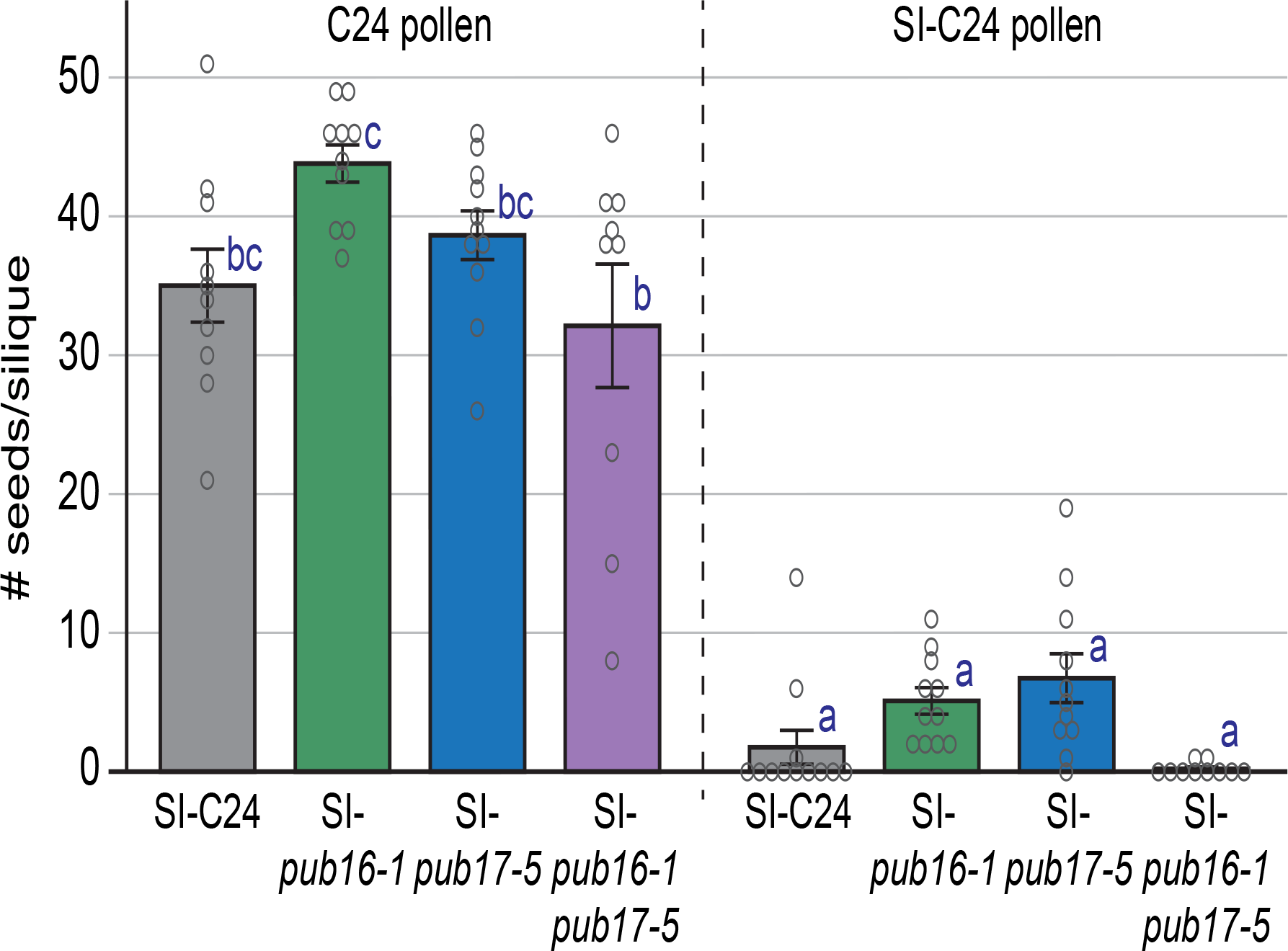
Seed yield for SI-C24 and SI-C24 *pub* mutant siliques. Pistils were pollinated with C24 pollen (left) or SI-C24 pollen (right), and siliques were harvested at 2-weeks post-pollination for counting. Data are shown as a bar graph of the means ± SE with all data points displayed. n = 10-11 siliques per line. Letters represent statistically significant groupings of P < 0.05 based on a one-way ANOVA with a Tukey-HSD post-hoc test.

## 3 DISCUSSION

The bulk of our understanding of the mechanisms underlying the Brassicaceae SI pathway originates from findings in *Brassica* species, and here we investigated if similar SI pathway components are regulating *Arabidopsis* SI. The factors for initiating SI pollen rejection, pollen SCR and stigma SRK, are conserved between *Brassica* and *Arabidopsis*, but investigations on the downstream players of this pathway suggest there are genus-specific differences (Abhinandan *et al*., 2022, Goring *et al*., 2023). For example, *A. thaliana* lost functional copies of the *SCR, SRK* and *ARC1* genes during the transition to selfing, and while functional copies of *SCR* and *SRK* are required to restore the SI trait in *A. thaliana*, ARC1 is not always required (Nasrallah *et al*., 2004, Boggs *et al*., 2009b, Indriolo and Goring, 2014, Indriolo *et al*., 2014, Fujii *et al*., 2020, Rozier *et al*., 2020). As well, introducing *SCR-SRK* pairs from closely-related *Arabidopsis* and *Capsella* SI species can reinstate the SI trait in transgenic *A. thaliana*, but *Brassica SCR-SRK* pairs fail to do so (Bi *et al*., 2000, Boggs *et al*., 2009a, Zhang *et al*., 2019, Yamamoto *et al*., 2022). These studies infer that at least some components of the downstream SI pathway have diverged between *Brassica* and *Arabidopsis*.

In this study, we investigated if the closely related *ARC1* paralogs, *PUB17* and *PUB16*, were functioning as an alternative to *ARC1* in the transgenic *A. thaliana* SI-C24 line. A number of PUBs have been implicated in plant immunity, often working together with redundant functions (Trujillo, 2018, Trenner *et al*., 2022). PUB17 was shown to be a positive regulator of plant immunity through the activation of programmed cell death (Yang *et al*., 2006). A putative K-homology RNA-binding protein, KH17, which acts as a negative regulator of immunity was proposed to be targeted by PUB17 for degradation (McLellan *et al*., 2020). For PUB16, litle is known other than its increased expression observed in flowers with GA treatment (Acosta *et al*., 2012). Previously we found a positive interaction between AlSRK_1_ and AlPUB17 (Indriolo and Goring, 2016), and our phylogenetic analysis showed that PUB17 and PUB16 are the two most closely related UND-PUBs to AlARC1 (Figure 1; Indriolo *et al*., 2012). The CRISPR/Cas9 system was used to generate *pub17* and *pub16* mutations in the C24 accession (Figure 2), and these mutations were then crossed into the transgenic *A. thaliana* SI-C24 line which carries the Al*SCR*_*b*_ and Al*SRK*_*b*_ transgenes and displays a strong SI phenotype (Iwano *et al*., 2015, Macgregor *et al*., 2022). Our experiments provided a thorough examination on the requirement of *PUB17* and *PUB16* at three different stages that are impacted by SI pollen rejection: pollen hydration, pollen tube growth, and seed yield. In response to SI-C24 pollen, the transgenic *A. thaliana* SI-C24 *pub* mutant pistils display strong SI pollen rejection responses with very low increases in pollen hydration, the absence of pollen tube growth in the pistil, and low seed counts (Figures 3-5). In all of these assays, there were no significant differences to the SI pollen rejection responses displayed by wildtype transgenic *A. thaliana* SI-C24 pistils. As well, the *pub17* and *pub16* mutant pistils did not have any impact on compatible pollen responses (Figures 3-5). Collecvtiely, these findings highlight that *PUB17* and *PUB16* are not compensating for the loss of *ARC1* in transgenic *A. thaliana* SI-C24 line, and do not influence the ability of SI-C24 line to reject SI pollen. Thus, based on this work, PUB17 and PUB16 are not required for self-pollen rejection in the transgenic *A. thaliana* SI-C24 line. In another study, *PUB2* which is more distantly related to *ARC1* (Figure 1) was tested in a transgenic *A. thaliana* Al*SCR*-Al*SRK* Col-0 line that has a weak SI trait and no role was found as well (Zhang *et al*., 2014).

In terms of cellular responses identified using transgenic *A. thaliana* SI lines, Iwano *et al*. (2015) discovered a rapid increase in cytosolic Ca^2+^ levels in the SI-C24 stigmatic papilla upon SI pollination. While a cytosolic Ca^2+^ flux was also observed with compatible pollen (Iwano *et al*., 2004) the size of the Ca^2+^ flux was much greater with SI pollen (Iwano *et al*., 2015) and would be predicted to disrupt actin cytoskeleton and vesicle secretion (Goring, 2017). Furthermore, Iwano *et al*. (2015) implicated Glutamate-Like Receptors (GLRs) to be responsible for the Ca^2+^ influx following SI pollination. Other cellular responses that have been identified in the transgenic *A. thaliana* SI lines include changes to actin filaments (Rozier *et al*., 2020) and the activation of autophagy (Safavian and Goring, 2013, Indriolo *et al*., 2014, Macgregor *et al*., 2022). We found that autophagy was necessary for the SI pathway as crossing autophagy mutants into transgenic *A. thaliana* SI-C24 and SI-Col-0 lines led to a partial breakdown of SI pollen rejection by the SI *atg* mutant pistils (Macgregor and Goring, 2022, Macgregor *et al*., 2022). Nevertheless, the mechanisms underlying the activation of autophagy and GLRs remain unknown. Thus, further investigations are required to uncover the signaling proteins directly activated by SRK and to determine how they are connected to the cellular events activated in the stigmatic papillae with SI pollen contact.

## 4 METHODS

### 4.1 Plant materials and growth conditions

All *A. thaliana* seeds (C24, SI-C24, and CRISPR/Cas9 deletion mutants) were sterilized and stratified at 4°C for 5 days in the dark. Stratified seeds were placed directly on Sunshine #1 soil supplemented with Plant Prod All Purpose 20-20-20 fertilizer for germination. Plants were grown in chambers under a 16-h light/8-h dark growth cycle at 22°C, with humidity kept under 50%. The *pub* mutants (*pub17-4, pub17-5, pub16-1* and *pub16-2*) generated through CRISPR/Cas9 (see below) were crossed with the transgenic *A. thaliana* SI-C24 line #15-1 (Iwano *et al*., 2015) to establish the single *pub* mutants in the SI-C24 background. The *pub16-1* mutant was then crossed with SI-C24 *pub17-5* to generate the SI-C24 *pub16-1 pub17-5* double mutant line. The SI-C24 line and *pub* mutants were genotyped by PCR using genotyping primers listed in Table S1.

### 4.2 Phylogenetic analysis

The amino acid sequences for *A. thaliana* PUB proteins with UND domains (Mudgil *et al*., 2004, Trenner *et al*., 2022) were retrieved from TAIR and used in BLAST searches of the *A. lyrata* genome in Phytozome to retrieve the *A*.*lyrata* UND-PUBs. In MEGA11, Muscle set at default parameters was used to align the amino acid sequences, and a maximum likelihood tree was built using default parameters and 1000 bootstrap replications. Gene identifiers for the genes used for this study are shown in Figure 1.

### 4.3 RT-PCR analysis of *PUB17* and *PUB16* expression in pistils

The top half of approximately 30 *A. thaliana* pistils (including the stigmas) were collected from stage 12 flower buds and flash frozen in liquid nitrogen. Total RNA was extracted using the standard protocol for the RNeasy Plant Mini kit (Qiagen). Extracted RNA was treated using RQ1 RNase-Free DNase (Promega) to eliminate genomic DNA contamination. cDNA was synthesized using Superscipt™ IV reverse transcriptase (ThermoFisher) with oligo(dT)20 primers and then used for PCR amplification (+RT). As a negative control, the cDNA synthesis reaction was setup with all components except the Superscipt™ IV reverse transcriptase and used for PCR amplification (-RT). Elongation Factor 1α2 (EF1A2; At1g07930) was used as a positive control for the RT-PCR reactions. See Table S1 for primers used.

### 4.4 Plasmid construction and plant transformation

The CRISPR/Cas9 vectors for creating deletion mutations in the C24 *PUB17* and *PUB16* genes were generated as previously described (Doucet *et al*., 2019a, Doucet *et al*., 2019b). The final pBEE401E vectors each contained two guide RNAs targeting the single exons of *PUB17* and *PUB16*. Wildtype *A. thaliana* C24 plants were transformed through *Agrobacterium*-mediated floral dip and to maximize transformation efficiency, plants were dipped twice, with a 7-day interval (Clough and Bent, 1998). Seeds from transformed plants were collected, stratified, and germinated on soil. 7-day old seedlings were sprayed with BASTA herbicide every 2 days, for a total of three times to select for transformants. Transformants were confirmed through detection of the BASTA resistance gene by PCR, and deletion mutations were identified through PCR genotyping. Plants in the T1 generation were self-fertilized, and in the T2 generation, two homozygous mutant alleles were detected for each gene *pub17-4, pub17-*5, *pub16-1*, and *pub16-2*. See Table S1 for primers used.

### 4.5 Pollen hydration, aniline blue staining, and seed set

All post-pollination assays used manual pollinations and were conducted as described in (Lee *et al*., 2020). All assays began with the emasculation of stage 12 flower buds from wildtype and mutant lines. Emasculated flower buds were wrapped with plastic wrap and stored in the growth chambers. To conduct the pollen hydration assay, the emasculated pistils were removed from the plant at 24-h post-emasculation and mounted on ½ MS plates. A single anther from SI-C24 open flowers was used to lightly pollinate pistils. Images were captured at 0- and 10-min post-pollination using a Nikon sMz800 microscope, and 10 pollen grains per pistil were randomly measured using the NIS-elements imaging software. This assay was repeated on 3 pistils per line, measuring a total of 30 pollen grains per group (*n* = 30). To conduct the aniline blue staining assay, emasculated pistils were pollinated with a single anther from SI-C24 open flowers at 24-h post-emasculation and were returned to the growth chambers. At 24-h post-pollination, the pistils were collected and stained as described in (Lee *et al*., 2020). Aniline blue-stained pistils were mounted on a microscope slide with sterile water, flattened with a coverslip, and imaged at 10X magnification on a Zeiss Axioskop2 Plus fluorescent microscope. Aniline blue images were captured using the UV laser, and brightfield images were captured using the built-in halogen lamp. The number of pollen tube/style were quanfitied for 8-16 pistils per line. To quantify seed yield, emasculated pistils were pollinated at 24-h post-emasculation using anthers from opened C24 or SI-C24 flowers and left for 10-14 days. Developed and matured siliques were collected and submerged in 70% (v/v) ethanol for 3-5 days or until siliques were cleared and seeds were visible (Beuder *et al*., 2020). Cleared siliques were mounted on a dry microscope slide with the septum facing upwards and imaged using a Nikon sMz800 microscope, and seeds were counted using the NIS-elements imaging software. For each line, the number of seeds/silique from 10 siliques were counted. For all statistical analyses, one-way analysis of variance (ANOVA) with Tukey’s honest significance difference (HSD) post-hoc tests were performed using SPSS (IBM, Armonk, NY, USAs).

## ACKNOWLEDGMENTS

We thank Samantha Lee, Erin Navaratnam, Cecilia Widjaja and Hamna Ammar for technical assistance, and members of the Goring lab for critically reading this article. We are also very grateful to Seiji Takayama (Nara Institute of Science and Technology; University of Tokyo) for providing the SI-C24 #15-1 seeds (*A. thaliana* C24). This research was supported by a grant from the Natural Sciences and Engineering Research Council of Canada to DRG.

## AUTHOR CONTRIBUTIONS

PB and DG designed the research; PB performed the research and wrote the paper; PB and DG analyzed data, edited the paper, and approved the final version.

## SUPPORTING INFORMATION

**Table S1.** List of primers used in this study.

