## Supplemental Table S1 for "Investigating a role for *PUB17* and *PUB16* in the self-incompatibility pathway in transgenic *Arabidopsis thaliana*"

**Table S1.** List of primers used in this study.

| Primer Name | Primer Sequence | Purpose of primer |
|---|---|---|
| 1F- C24 pub17-4 target site 1 FP | ATATATGGTCTCGATTGGGAAGCGCAAGAAAATG CAGGTTTTAGAGCTAGAAATAGCAAGTTAAAAT | gRNAs used for generating *pub17-4* deletion in C24 |
| 3R- C24 pub17-4 target site 2 RP | ATTATTGGTCTCTAAACCCGAAGAGATTGAAGCTA GGCAATCTCTTAGTCGACTCTACCAATA | |
| 3F- C24 pub17-5 target site 1 FP | ATATATGGTCTCGATTGCCTAGCTTCAATCTCTTC GGGTTTTAGAGCTAGAAATAGCAAGTTAAAAT | gRNAs used for generating *pub17-5* deletion in C24 |
| 2R- C24 pub17-5 target site 2 RP | ATTATTGGTCTCTAAACCTGCTACACCTTCGTTCT TTCAATCTCTTAGTCGACTCTACCAATA | |
| PUB16 G5 | ATTATTGGTCTCTAAACCGTGGGAGGCTAAGGCT AACCAATCTCTTAGTCGACTCTACCAATA | gRNAs used for generating *pub16-1* deletion in C24 |
| PUB16 G6 | ATATATGGTCTCGATTGTTGTATCGGTGACTTTAC GAGTTTTAGAGCTAGAAATAGCAAGTTA | gRNAs used for generating *pub16-1* deletion in C24 |
| PUB16 G5 | ATTATTGGTCTCTAAACCGTGGGAGGCTAAGGCT AACCAATCTCTTAGTCGACTCTACCAATA | gRNAs used for generating *pub16-2* deletion in C24 |
| PUB16 G8 | ATATATGGTCTCGATTGTAGCAGAATCGGATAAAC CAGTTTTAGAGCTAGAAATAGCAAGTTAAAAT | gRNAs used for generating *pub16-2* deletion in C24 |
| BASTA FP | AAGCACGGTCAACTTCCGTA | Genotyping for BASTA resistance gene |
| BASTA RP | GAAGTCCAGCTGCCAGAAAC | |
| SCRb FP | ATGAGGAATGCTACTTTCTTC | Genotyping SI-C24 for *Al-SCRb* (Iwano et al. 2015) |
| SCRb RP | TAGCAAAATCTACAGTCGCATA | |
| SRKb FP | ACCAAGATTCACGGTTCAGG | Genotyping SI-C24 for *Al-SRKb* (Iwano et al. 2015) |
| SRKb RP | ACGCTGTTCATGTGTCGAAG | |
| PUB17 KO FP | TTAGCCCCCGTTGACCTTTC | Genotyping for CRISPR/Cas9 *PUB17* deletion |
| PUB17 KO RP | AATCGCAGGTGCTCTCAACA | |
| PUB17 INT FP | GAGCCAACGGGATCGGTTAT | Genotyping for CRISPR/Cas9 *PUB17* internal |
| PUB17 INT RP | TTTGTTCGCTTCAACCGCAG | |
| PUB16 KO FP | AATCCGCTCATCTCTGCTTC | Genotyping for CRISPR/Cas9 *PUB16* deletion |
| PUB16 KO RP | TGTGGTATCCGCTCCTTCTC | |
| PUB16 INT FP | TCCGAAAACAGGTCAAGTCC | Genotyping for CRISPR/Cas9 *PUB16* internal |
| PUB16 INT RP | AACGAACCAGCTTGGGTATG | |
| PUB17 LP | GAGCCAACGGGATCGGTTAT | PUB17 RT-PCR primers |
| PUB17 RP | TTTGTTCGCTTCAACCGCAG | |
| PUB16 LP | TCCGAAAACAGGTCAAGTCC | PUB16 RT-PCR primers |
| PUB16 RP | AACGAACCAGCTTGGGTATG | |
| EF1A2 LP | GCTCCTGGTCATCGTGATTT | Elongation Factor 1α2 (EF1A2, At1g07930) RT-PCR primers |
| EF1A2 RP | CAGTCAAGGTTGGTGGACCT | |
